# Genetic variability in response to Aβ deposition influences Alzheimer’s risk

**DOI:** 10.1101/437657

**Authors:** Dervis A. Salih, Sevinc Bayram, Manuel S. Guelfi, Regina Reynolds, Maryam Shoai, Mina Ryten, Jonathan Brenton, David Zhang, Mar Matarin, Juan Botia, Runil Shah, Keeley Brookes, Tamar Guetta-Baranes, Kevin Morgan, Eftychia Bellou, Damian M. Cummings, John Hardy, Frances A. Edwards, Valentina Escott-Price

## Abstract

Genetic analysis of late-onset Alzheimer’s disease risk has previously identified a network of largely microglial genes that form a transcriptional network. In transgenic mouse models of amyloid deposition we have previously shown that the expression of many of the mouse orthologs of these genes are co-ordinately up-regulated by amyloid deposition. Here we investigate whether systematic analysis of other members of this mouse amyloid-responsive network predicts other Alzheimer’s risk loci. This statistical comparison of the mouse amyloid-response network with Alzheimer’s disease genome-wide association studies identifies 5 other genetic risk loci for the disease (*OAS1, CXCL10, LAPTM5, ITGAM* and *LILRB4*). This work suggests that genetic variability in the microglial response to amyloid deposition is a major determinant for Alzheimer’s risk.

**One Sentence Summary:** Identification of 5 new risk loci for Alzheimer’s by statistical comparison of mouse Aβ microglial response with gene-based SNPs from human GWAS

## Main Text

All the mutations in the genes causing early-onset Alzheimer’s disease (AD) alter APP processing such that amyloid deposition becomes more likely (*1*). In contrast, with the exception of some rare variants in APP processing enzymes (*2–5*), the majority of the risk in late-onset disease has been shown to be due to sequence variability in genes expressed in the innate immune system (largely microglial) and lipid metabolism (*6*). When we identified the microglial gene *TREM2* (*7*) as a potent risk gene for late-onset disease, we confirmed earlier reports that its expression was strongly increased by amyloid deposition in *APP* transgenic mice (*7–10*). In a genome-wide expression study of transgenic *APP/PSEN1* mice during pathology development, we noted that *Trem2* was one of the genes whose expression was up-regulated the most in relation to amyloid deposition and that *Trem2* expression showed a strong correlation with an entire network of genes co-expressed in the innate immune system. This immune module of genes showed a remarkable correlation to amyloid pathology and contained orthologs of other established Alzheimer’s risk genes such as *Abca7* and *Ms4a6d* (correlation = 0.87; p = 6e^−32^)(*9, 11*). Notably, the two AD risk loci for *ABI3* and *PLCG2* identified subsequent to our study were also present in this network (*12*), suggesting that this amyloid-responsive immune network may predict future risk genes for AD.

An important outstanding question is whether late-onset AD is mostly due to an inadequate cellular response to rising Aβ and its deposition, particularly due to sequence and expression variability in genes expressed by the innate immune system and/or involved in lipid processing. This hypothesis is difficult to study in human post-mortem tissue because after an extended period of disease the proportion of cell types in the brain have changed and the remaining cells show extensive compensatory changes in gene expression. With these questions in mind, we determined whether surveying the gene expression network that responds robustly to amyloid pathology could be used to identify further AD risk loci. Although amyloid mouse models have clear limitations in that they do not show tau tangles or neuronal loss, they allow us to study the time-course response of a healthy innate immune system reacting to Aβ, in which the innate immune cells have the ability to ultimately prevent Aβ killing neurons. Our previous expression network was constructed using expression arrays (*9*). Because these microarrays are limited by their probe content and have a limited dynamic range, we have now sequenced the transcriptome using RNA-seq and reconstructed a higher resolution expression network. The new full microglial module of genes shows a dramatic correlation with Aβ pathology (correlation = 0.94; p < 3e^−41^), and contains the mouse orthologs of existing GWAS loci *TREM2*, *ABI3*, *CD33*, *INPP5D*, *MS4A6D*, *SPI1*, *PLCG2*, *RIN3, HLA* and *APOE* (Table S1). The genes showing the tightest expression correlation/Aβ-response within the module form the network shown in Fig. 1 and Table S2 (top 147 genes from a total of 1,584 genes with up-regulated expression as part of the immune module based on the topological overlap measure, TOM, see Methods). This network is broadly similar to the network derived from the analysis of the same RNA by microarray methods (*9*), and importantly closely resembles microglial networks published by other groups using different amyloid mouse models (*13–17*), suggesting this is a conserved network of genes that can be reliably identified using different methodologies. *Trem2* forms a hub gene in our network indicating that *Trem2* expression is highly correlated to many other genes in the network, and may drive the expression response of this network. In line with this idea, *Trem2* has been shown to regulate at least part of this immune module (*13, 14, 16*). The network we identified also is broadly similar to a human network of immune genes containing *TYROBP*, *TREM2*, *MS4A* family genes, *C1Q* members and *CD33*, identified from human pathology tissue bearing in mind the caveats discussed above (*18, 19*), suggesting this mouse Aβ-response gene network behaves similarly in humans.

**Fig. 1.**
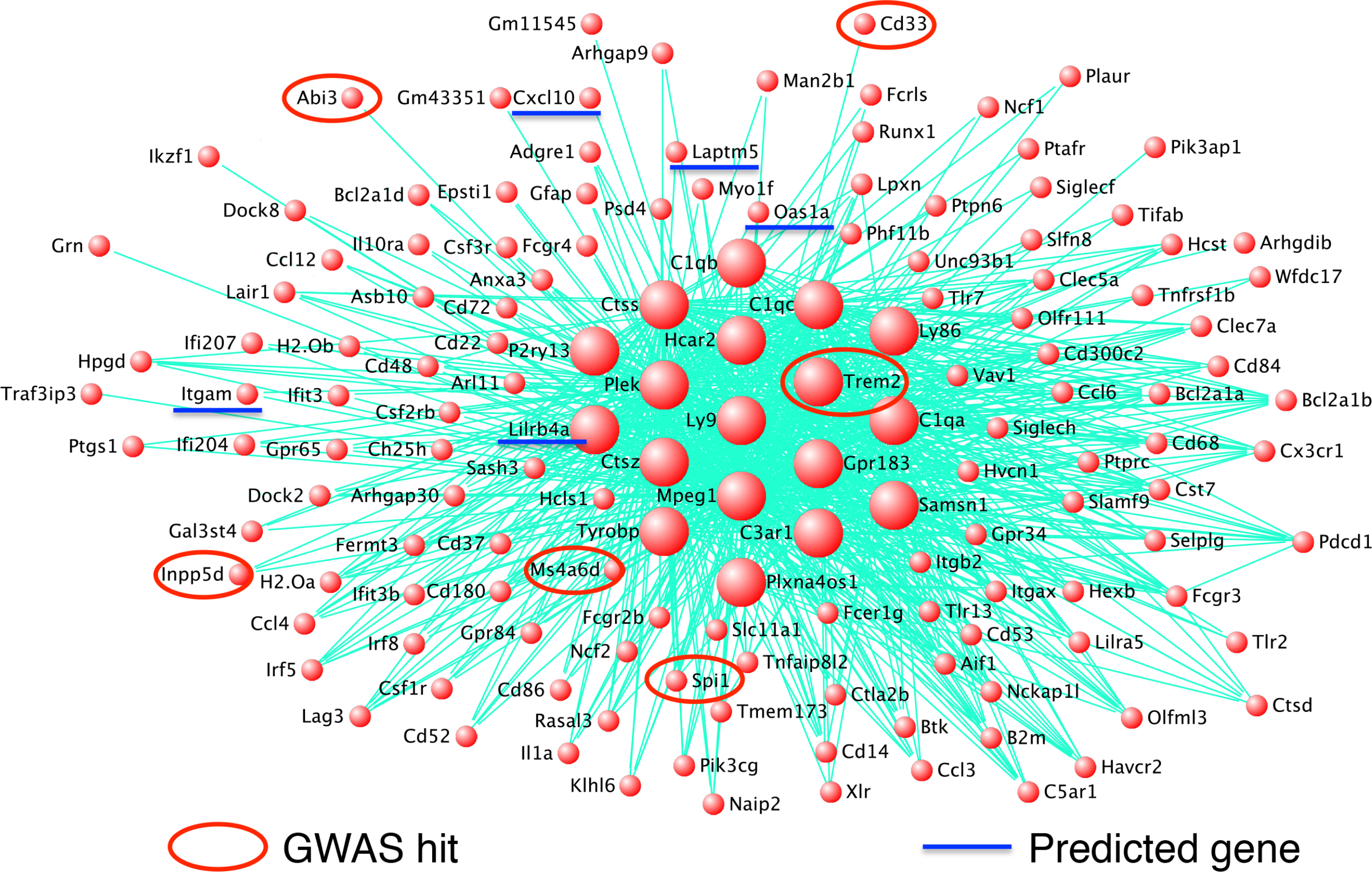
Amyloid-responsive immune network of genes featuring several orthologs of established GWAS variants associated with AD, predicts the importance of five new genes that may influence the risk of developing AD. Network plot using VisANT reveals key drivers of an immune module from RNA-seq derived gene expression from the hippocampus of wild-type and amyloid mice. Red circles show established GWAS genes associated with AD including *Trem2*, *Cd33*, *Abi3* and *Spi1*. Blue underline shows genes predicted to confer increased risk of AD by overlapping strongly amyloid-responsive gene expression data in amyloid mice with analyses identifying combinations of adjacent human SNPs within individual genes showing significant associations with AD (see Methods, *20*). Genes shown in this network display up-regulated expression in response to amyloid deposition. Larger red spheres represent ‘hub genes,’ those showing the greatest number of connections to other genes in the network, and include *Trem2*, *Tryobp*, *Lilrb4a*, *P2ry13*, *Ctss*, *Ctsz*, *Mpeg1* and *Plek*, which are likely to play important roles in driving microglial function.

Within our mouse immune network, we first confirmed that several members were orthologs of AD loci variants using the data from the Alzheimer’s disease genetic consortium (*11, 20*)(Table 1). We then asked whether the other members of the mouse microglial amyloid-response network overlapped with individual human genes containing multiple SNPs associated with AD by cross-referencing gene-based statistical approaches (*20*). Overall, we found there was an enrichment of human genes with significant AD-associated SNPs within this amyloid-responsive network. This enrichment was more than would be expected by chance alone, even after the established GWAS loci were excluded (p = 1.91×10^−5^ for highly connected network genes, Fig. 1, top 147 genes, versus p = 7.32×10^−4^for the entire module of 1,584 genes, Table S1). As a comparison to the mouse amyloid-responsive network, the mouse tau-responsive immune network was not significantly enriched for human genes with AD-associated SNPs when the central portion of the tau network containing the highly connected genes were considered, after the established GWAS loci were excluded as before (p = 0.92), although *Apoe* is part of this module (Fig. S1, top 137 genes from a total of 2,299 genes in the immune module based on the TOM). When the entire module of tau-responsive immune genes (2,299 genes) was considered there was a significant enrichment, p = 4.63×10^−6^, suggesting that a proportion of AD-associated SNPs appear in microglial genes that have mouse orthologs, but are less responsive to tau pathology compared to amyloid pathology. The amyloid network analyses identified 5 genes within the mouse microglial network whose human orthologes contained SNPs significantly associated with AD, counting the genes within 0.5 Mb as one locus (see Methods, *20*). These 5 genes, *OAS1*, *CXCL10*, *LAPTM5*, *ITGAM* and *LILRB4*, have not been previously reported as having variants significantly associated with AD using traditional GWAS approaches (Table 1, Fig. S2-4). Indeed the amyloid-responsive sub-network of these 5 novel genes with the established GWAS loci *TREM2*, *ABI3*, *CD33*, *INPP5D, SPI1* and *MS4A6D* (Fig. 1) is not highly connected in an innate immune gene network associated with tau pathology (Fig. S1), suggesting this sub-network is more responsive to amyloid pathology than other pathologies. Furthermore, in common with the existing 6 known GWAS-associated genes, the 5 novel genes we identify respond very early to Aβ deposition, with gene expression increasing from 4 months of age in the homozygous *APP/PSEN1* mice (Fig. S5).

**Table 1.** The genes predicted to contain SNP variants associated with AD together with established loci associated with AD from GWAS. Genes predicted to confer increased risk of AD by overlapping strongly amyloid-responsive gene expression data in amyloid mice (Fig. 1) with analyses identifying combinations of adjacent human SNPs within individual genes showing significant associations with AD (see Methods; *20*). The SNP positions are provided for build 37, assembly Hg19, as in IGAP study (*11*). The SNP with the most significant p-value within each gene is denoted as ‘Best SNP,’ from the IGAP stage 1 dataset. The effect size (coefficient of the logistic regression) is provided for the best reported SNP from IGAP data; a positive number indicates that the allele increases risk of AD, and so a negative number indicates the allele is protective. The allele frequency from the IGAP study is also provided. The established genes altering risk for AD from GWAS are given for comparison.

| Mouse symbol (MGI) | Human symbol (HGNC) | NCBI ID | Human Chromosome | Start Location | End Location | Number of SNPs | Gene p-value (adj for GC) | Best SNP | Best SNP Location | Best SNP p-value | Effect size | Risk Allele | Frequency |
| --- | --- | --- | --- | --- | --- | --- | --- | --- | --- | --- | --- | --- | --- |
| ***Predicted genes*** |  |  |  |  |  |  |  |  |  |  |  |  |  |
| *Laptm5* | *LAPTM5* | 7805 | 1 | 31205315 | 31230683 | 45 | 0.00285 | rs1623695 | 31210852 | 0.000764 | -0.0817 | T | 0.2065 |
| *Cxcl10* | *CXCL10* | 3627 | 4 | 76942271 | 76944650 | 6 | 0.00227 | rs8878 | 76942300 | 0.00144 | 0.0508 | A | 0.4657 |
| *Oas1a* | *OAS1* | 4938 | 12 | 113344739 | 113357712 | 17 | 0.000388 | rs1131454 | 113348870 | 3.92E-05 | 0.1004 | A | 0.5655 |
| *Itgam* | *ITGAM* | 3684 | 16 | 31271288 | 31344213 | 151 | 0.00571 | rs9928397 | 31320901 | 0.000671 | 0.1079 | T | 0.0894 |
| *Lilrb4a* | *LILRB4* | 11006 | 19 | 55174124 | 55179848 | 22 | 0.00666 | rs731170 | 55176262 | 0.000272 | 0.0683 | A | 0.3054 |
| ***Established GWAS genes*** |  |  |  |  |  |  |  |  |  |  |  |  |  |
| *H2-Ob* | *HLA-DOB* | 3112 | 6 | 32780540 | 32784825 | 35 | 0.00354 | rs2070121 | 32781554 | 0.00138 | 0.0931 | A | 0.0796 |
| *Trem2* | *TREM2* | 54209 | 6 | 41126246 | 41130922 | 1 | 0.00258 | rs7748513 | 41127972 | 0.00182 | -0.1293 | A | 0.9633 |
| *Spi1* | *SPI1* | 6688 | 11 | 47376409 | 47400127 | 47 | 1.34E-06 | rs10437655 | 47391948 | 1.99E-06 | 0.0759 | A | 0.4037 |
| *Ms4a6d* | *MS4A6A* | 64231 | 11 | 59939080 | 59950674 | 22 | 3.07E-10 | rs7935829 | 59942815 | 1.64E-10 | 0.1011 | A | 0.5989 |
| *Abi3* | *ABI3* | 51225 | 17 | 47287589 | 47300587 | 37 | 0.00228 | rs2158512 | 47290253 | 9.22E-07 | 0.154 | T | 0.726 |
| *Cd33* | *CD33* | 945 | 19 | 51728335 | 51743274 | 23 | 1.95E-06 | rs12459419 | 51728477 | 6.49E-08 | -0.0945 | T | 0.3102 |

Aspects of the amyloid-responsive network we identify in our analysis containing the 5 new genes with the existing 6 GWAS loci are broadly similar to microglial networks we and others have previously identified in human brain analyses. Zhang and colleagues identified an AD-relevant network centered on *TYROBP* and *TREM2* which contained *ITGAM* and *LAPTM5* (*18*) and we described a human microglial network containing *LAPTM5*, *ITGAM* and *LILRB4* (*19*). We then determined whether these novel Alzheimer’s risk loci, derived from a mouse Aβ-response network were present in independent datasets of human brain co-expression networks. Cross referencing the network (see Methods) with the data from the ROS/MAP project (*21, 22*), and BRAINEAC (*23*) datasets revealed that *LAPTM5*, *ITGAM* and *LILRB4* clustered together in the same network in the ROSMAP based co-expression networks, together with many of the GWAS risk genes for AD, and with *SPI1*, the myeloid cell transcription factor (*24*)(Fig. S6; Fisher’s Exact test Bonferroni corrected p = 1.34×10^−13^ for AD). We confirmed these module memberships in the BRAINEAC data for control brains generated in our own lab and found essentially the same results (data not shown). Interestingly, we found that SPI1 was bound to the regulatory regions of *Laptm5* and *Itgam*, along with binding to established AD risk gene orthologs *Trem2*, *Abi3*, *Inpp5d*, *Ms4a6d* and *Spi1* itself, by searching data from a chromatin immunoprecipitation experiment against SPI1 in mouse microglial-like BV-2 cells (*25*). This finding was supported by mining for regulatory features and *cis*-regulatory modules in the amyloid-response network genes using i-*cis*Target that uses a vast library of regulatory data (*26*). Together, these findings suggest that a number of the predicted and established AD risk genes may be regulated by SPI1, which itself alters AD risk by coordinating a program of microglial-expressed genes (*24*).

Since most GWAS loci are thought to operate by regulating the expression of neighboring genes (*24, 27, 28*), for each of the 5 potential AD-associated genes we performed a colocalisation analysis to test the association between AD loci located within these genes and loci regulating these genes’ expression (eQTLs; (*29*). eQTLs were obtained from two previously published datasets using baseline and stimulated human-derived monocytes and iPSC-derived macrophages (*30, 31*). In these studies, macrophages and monocytes were stimulated with various immunostimulants to activate distinct, well-characterised immune signaling pathways, including those broadly associated with bacterial and viral responses. Interestingly, we identified 3 colocalisations between AD loci and eQTLs regulating *OAS1* gene expression, all of which were identified in stimulated states, suggesting that this association is only active in certain environmental conditions (Fig. 2 and Fig. S7-8), in particular those designed to model monocyte/macrophage priming or more chronic inflammation.

**Fig. 2.**
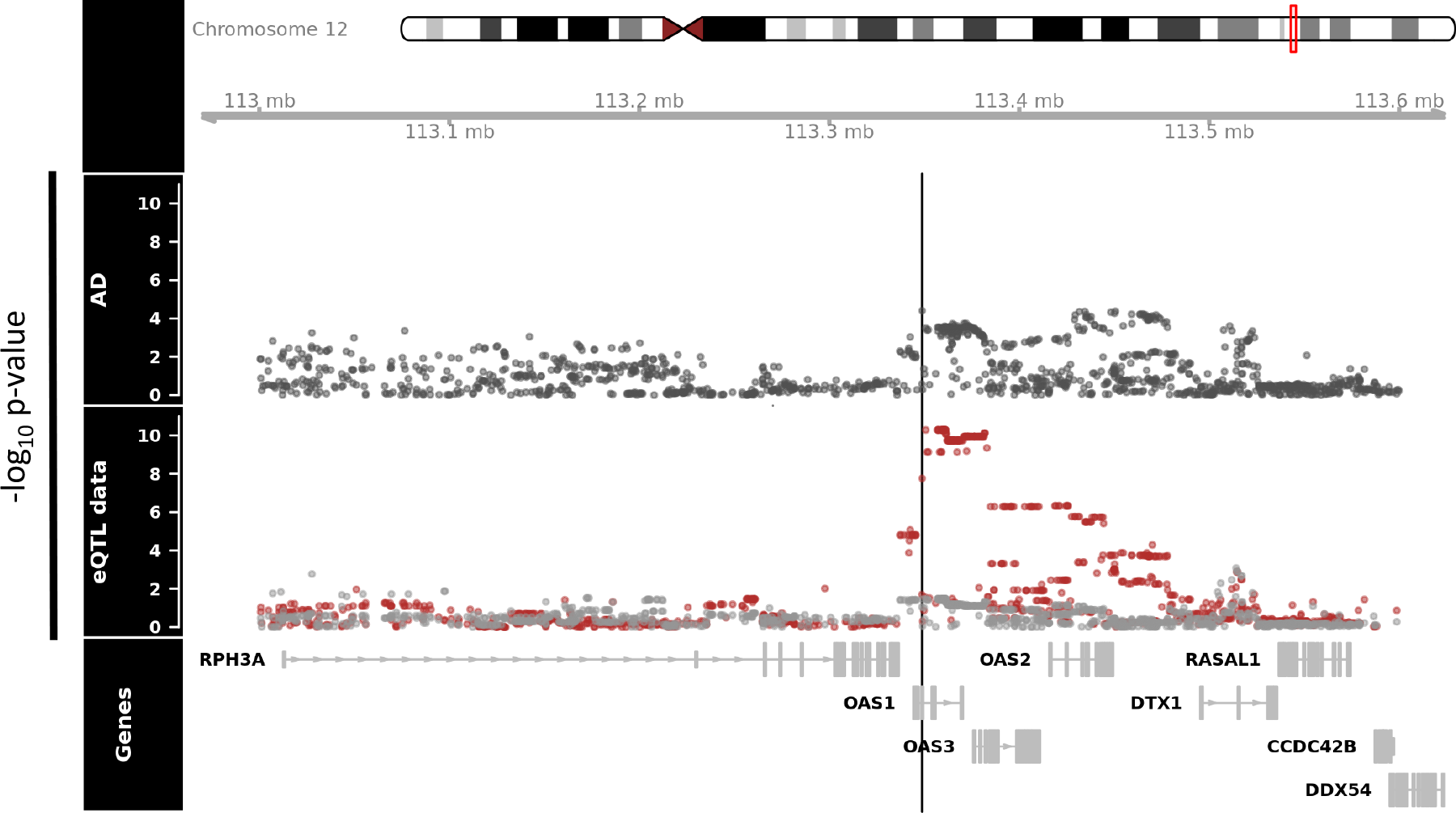
Colocalisation of AD GWAS loci with eQTLs derived from baseline and stimulated iPSC-derived macrophages. Colocalisation of AD loci and eQTLs targeting *OAS1* in baseline and stimulated states (IFNγ and Salmonella, 18 and 5 hours respectively). In the eQTL panels, grey and red data points represent macrophages at baseline or stimulated with both IFNγ and Salmonella, respectively. The best AD locus in *OAS1*, rs1131454 (p-value = 3.92 × 10-5), is highlighted with the black line. IFNγ, interferon-γ. Numerical results are reported in Table S3.

Surveying the literature on our genes of interest revealed that *OAS1* (2-prime,5-prime oligoadenylate synthetase 1) is involved in the regulation of cytokine expression (*32*). *OAS1* is induced by interferons (*33*), which supports our eQTL analysis showing that the best SNP we identified for *OAS1* appears in a locus which acts as an eQTL in response to interferon-γ (IFNγ; Fig. 2 and Fig. S7-8). *OAS1* can additionally activate Ribonuclease L which degrades viral RNA and inhibits viral replication (*33*). *CXCL10* (IP-10; chemokine, CXC motif, ligand 10) is a proinflammatory cytokine that has been reported to have increased concentrations in the AD-brain, particularly associated with amyloid plaques (*34*), and *CXCL10* increases plaque pathology in *APP/PSEN1* transgenic mice (*35*). *CXCL10* was found to increase in older people and in AD, and correlated with cognitive decline (*36*). *LAPTM5* (lysosome-associated protein, transmembrane 5) is associated with amyloid pathology in transgenic mice (*17*), and *LILRB4* (leukocyte immunoglobulin-like receptor, subfamily B, member 4), has also been shown to be increased with amyloid pathology and specifically associated with amyloid plaques (*15, 37, 38*). The functions of *LAPTM5* and *LILRB4* have not been well characterized, but are thought to suppress the activation of a variety of immune cells. *ITGAM* (alpha-chain subunit of the heterodimeric integrin complement receptor alpha-M-beta-2, also known as CD11b or CR3A), is a cell surface receptor involved in activation, migration and phagocytosis of immune cells, so much so that ITGAM is used as a marker of activated microglia (*37, 39, 40*), and is involved in systemic lupus (*41*). *ITGAM* was highlighted in recent genetic and functional analyses as being a likely AD risk gene, whose expression was driven by SPI1, and related to amyloid pathology in mice and humans (*17, 18, 24, 37, 42, 43*).

The importance of this work is two fold. First, by identifying more genetic loci involved in pathogenesis, we derive a more complete insight into the cellular processes and molecular mechanisms underlying the disease. In this regard this work is complementary to that of Huang and colleagues (*24*), showing that microglial SPI1-driven transcription is a common feature of many Alzheimer’s loci. These findings are also consistent with previous work on *Trem2* (*8, 13, 14, 16, 44*) and *CD33* (*27, 28*) suggesting these risk genes are crucial in controlling the microglial response to amyloid-induced damage. Understanding the mechanisms of function of TREM2 and the amyloid-responsive sub-network identified here may be useful for leveraging therapeutic opportunities. Second, and perhaps of greater importance, this work implies that, overall, how well an individual responds to amyloid deposition at the cellular and gene expression level plays a large part in determining ones risk of disease, and this may be used to predict the chances of developing AD before irreversible neurodegeneration sets in.

### URLs of databases used

Mouseac: www.mouseac.org

Braineac: www.braineac.org

1,000 genomes: http://www.1000genomes.org/ and http://www.internationalgenome.org/

Coloc: https://github.com/chr1swallace/coloc

Bioconductor: https://bioconductor.org/biocLite.R

Coloc: https://github.com/chr1swallace/coloc

Bioconductor: https://bioconductor.org/biocLite.R

ROS/MAP: https://www.synapse.org/#!Synapse:syn3219045

i-*Cis*Target: https://gbiomed.kuleuven.be/apps/lcb/i-cisTarget/

GTEx V6 gene expression: https://gtexportal.org/home/

Coexp: https://github.com/juanbot/coexp

## Methods

### Mouse and transcriptome work

Total RNA was used from the same mice as described in Matarin *et al.* (*9*). The quality and concentration of the total RNA was assessed using capillary electrophoresis of each sample. RNA-seq library preparation and sequencing was performed by Eurofins Genomics (strand-specific cDNA libraries with polyA selection), by Illumina (HiSeq 2500) sequencing (2× 100 bp paired-end; multiplex 12 samples per lane - 28M reads). Adaptors and low quality base pairs were removed from FASTQ files using Trim Galore (Babraham Bioinformatics).

Transcripts were quantified with Salmon (*45*), using gene annotation from ENSEMBL GRCm38. Salmon was used because it incorporates GC correction and accounts for fragment positional bias. To get gene level quantification from the transcripts, and correct for average transcript length and library size, expressed as transcripts per million (TPM), the tximport R package was used (*46*). TPM values were log2 transformed, and genes were considered expressed when log2 TPM values displayed a mean >1.5 for a given gene for at least one group of mice, when gene TPM values were averaged for each genotype at each age (resulting in a total of 18,562 genes expressed).

Weighted co-expression network analyses (WGCNAs) was performed as described in Matarin *et al.* (*9*). Coexpression networks were built using the WGCNA package in R. Genes with variable expression patterns (coefficient of variation >5% for wild-type and amyloid mice, or wild-type and tau mice) from normalized log2 TPM values were selected for network analyses resulting in 13,536 genes for network analyses (*47–50*). The module of genes with the highest significant correlation with amyloid or tau pathology was selected for analysis (amyloid, correlation 0.94, p = 3e-41; tau, correlation 0.82, p =4e-12). TOM connectivity values were used to plot the network diagrams (TOM > 0.39 for amyloid-responsive module, and TOM > 0.36 for tau-responsive module). Hub genes were considered to be those with at least 15 connections to other genes.

### Genetic Analysis

The lists of mouse genes were converted to the lists of human genes using convertMouseGeneList() function, library biomaRt in R downloaded from https://bioconductor.org/biocLite.R.

The significance of the association of human genes to AD was assessed as described in (*20*). Briefly, the IGAP (*11*) summary statistics calculated for each SNP in a sample of 17,008 AD cases and 37,154 controls were used to derive the gene-based p-values. SNPs were assigned to genes if they were located within the genomic sequence lying between the start of the first and the end of the last exon of any transcript corresponding to that gene. The chromosome and location for all currently known human SNPs along with their assignment to genes were taken from the dbSNP132 database (build 37.1). If a SNP belongs to more than one gene, it was assigned to each of these genes. Data from the 1,000 genomes project (release Dec2010) were used as a reference panel for both (a) SNP imputation, and (b) calculation of LD between markers (*51*). An approximate statistical approach (*52*) which controls for LD and different number of markers per gene, was used to derive the gene-based p-values. Prior to the gene-based analyses all individual SNP p-values were corrected for genomic control.

We calculated the significance of the excess *number* of genes attaining the specified thresholds (0.05, 0.01 and 0.001) based upon the assumption that, under the null hypothesis of no association, the number of significant genes at a significance level of *α* in a scan is distributed as a binomial (*N, α*), where *N* is the total number of genes, assuming that genes are independent. Genes within 0.5Mb of each other are counted as one signal when calculating the observed number of *significant* genes. This prevents significance being inflated by LD between genes, where a single association signal gives rise to several significantly-associated genes. The over-representation p-value was calculated using a Z-test comparing the number of observed independent significant genes with the expected number of significant genes with corresponding variance (=*N*\**a* *(1−*a*), where *N* is the total number of independent genes in the network, and *a* is the significance threshold). We report the genes at the gene-based p-value threshold 0.01, where the excess of observed significant genes was the highest.

### Human sample co-expression network construction and annotation

We generated co-expression networks from RNA-seq based gene expression profiling of 635 pre-frontal cortex samples from the ROS/MAP project (*21, 22, 53*). We used cognitive decline as a covariate to construct four networks: all samples network, not AD, probable AD and AD. We used WGCNA (*50*) with an optimization for constructing more biologically meaningful co-expression networks (*54*). We corrected for batch effects using ComBAT (*55*), obtained unknown hidden effect covariates with SVA (*56*), and used the residuals obtained by regressing the gene expression with SVA covariates, age and gender.

Then we annotated the network modules for enrichment of Gene Ontology, REACTOME (*57*), and KEGG (*58*) pathways using gProfileR (*59*).

### Colocalisation with monocyte eQTL data sets

We applied coloc (version 3.1, see URLs) to test for colocalisation between AD loci surrounding the five novel identified genes (*OAS1*, *CXCL10*, *LAPTM5*, *ITGAM*, and *LILRB4*) and eQTLs (*29*). While no microglial eQTL datasets exist to date, eQTL analyses have been performed using monocytes and iPSC-derived macrophages (at rest and stimulated with various immunostimulants, such as IFN-γ)(*30, 31*). We ran coloc using default parameters and priors on all SNPs that: 1) had eQTLs tagging one of the 5 novel genes (this included all tested SNP-gene associations, including non-significant eQTLs); and 2) had overlapping SNPs in the AD GWAS. We excluded all loci in which PP3 + PP4 < 0.8, to exclude loci where we were underpowered to detect colocalisation. Loci with PP4/PP3 ≥ 2 were considered colocalised due to a single shared causal variant (PP4), as opposed to two distinct causal variants (PP3).

## Acknowledgements

DAS and FAE are funded by ARUK. JH and VEP are members of the UKDRI. JH and FAE are supported by the Cure Alzheimer’s Fund, JH is supported by the Dolby Foundation. The University of Nottingham Group is funded by ARUK and hosts the ARUK Consortium DNA Bank, with the following members: Tulsi Patel^1^, David M. Mann^2^, Peter Passmore^3^, David Craig^3^, Janet Johnston^3^, Bernadette McGuinness^3^, Stephen Todd^3^, Reinhard Heun^3^, Heike Kölsch^5^, Patrick G. Kehoe^6^, Emma R.L.C. Vardy^7^, Nigel M. Hooper^2^, Stuart Pickering-Brown^2^, Julie Snowden^8^, Anna Richardson^8^, Matt Jones^8^, David Neary^8^, Jenny Harris^8^, A. David Smith^9^, Gordon Wilcock^9^, Donald Warden^9^ and Clive Holmes^10^

^1^Schools of Life Sciences and Medicine, University of Nottingham, Nottingham NG7 2UH, UK, ^2^Institute of Brain, Behaviour and Mental Health, Faculty of Medical and Human Sciences, University of Manchester, Manchester M13 9PT, UK, ^3^Centre for Public Health, School of Medicine, Queen’s University Belfast, BT9 7BL, UK, ^4^Royal Derby Hospital, Derby DE22 3WQ, UK ^5^Department of Psychiatry, University of Bonn, Bonn 53105, Germany, ^6^School of Clinical Sciences, John James Laboratories, University of Bristol, Bristol BS16 1LE, UK, ^7^Salford Royal NHS Foundation Trust, ^8^Cerebral Function Unit, Greater Manchester Neurosciences Centre, Salford Royal Hospital, Stott Lane, Salford M6 8HD, UK, ^9^University of Oxford (OPTIMA), Oxford OX3 9DU, UK ^10^Clinical and Experimental Science, University of Southampton, Southampton SO17 1BJ, UK.

